# Lyophilized Cell-Secreted Matrix as a Bioactive Substrate for Chondrocyte Expansion and Redifferentiation

**DOI:** 10.64898/2026.02.13.705752

**Authors:** Mira Hammad, Benoît Domin, Alexis Veyssière, Benoît Bernay, Catherine Baugé, Karim Boumédiene

## Abstract

Articular cartilage repair is limited by the poor regenerative capacity of chondrocytes and their rapid dedifferentiation during in vitro expansion. This study investigated whether a decellularized and lyophilized cell-secreted matrix (CSM) could function as a bioactive material to regulate cell behavior, promote chondrogenic differentiation, and attenuate or reverse chondrocyte dedifferentiation without exogenous growth factor supplementation. CSM was generated from rabbit auricular perichondrial cells, decellularized, lyophilized, and characterized by histology, biochemical assays, and proteomic analysis. The resulting matrix was enriched in structurally and functionally relevant extracellular matrix proteins, including collagens, fibronectin, fibrillin, proteoglycans, and matricellular regulators, with minimal intracellular contamination and good batch-to-batch reproducibility. Functionally, CSM supported robust adhesion and proliferation of allogeneic and xenogeneic cells. Human articular chondrocytes cultured on CSM exhibited enhanced proliferation, sustained expression of cartilage-specific markers, and preserved type II collagen production over serial passages compared with standard plastic culture. CSM also promoted chondrogenic differentiation of human progenitor cells and partially reversed established chondrocyte dedifferentiation, as evidenced by increased expression of COL2A1, ACAN, SOX9, and COMP, with reduced COL1 expression and no induction of hypertrophic markers. These findings demonstrate that lyophilized CSM is a stable, off-the-shelf biomaterial capable of directing chondrocyte fate through intrinsic matrix-derived cues, highlighting its potential for cartilage tissue engineering and cell manufacturing applications.

## Introduction

Articular cartilage repair remains a major clinical challenge in orthopedic surgery due to the intrinsic avascularity and limited regenerative capacity of cartilage tissue. Cell-based therapeutic strategies, including matrix-assisted autologous chondrocyte transplantation (MACT), are widely employed for the treatment of focal cartilage defects and have demonstrated encouraging clinical outcomes [1]. However, the success of these approaches critically depends on the availability of large numbers of phenotypically stable chondrocytes. In vitro expansion through serial passaging, which is required to generate clinically relevant cell numbers, is strongly associated with chondrocyte dedifferentiation and progressive loss of cartilage-specific functionality [2].

Chondrocyte dedifferentiation is characterized by profound changes in extracellular matrix (ECM) synthesis and cellular metabolism, driven by an imbalance between anabolic and catabolic signaling pathways. This process is marked by a downregulation of cartilage-specific markers, particularly type II collagen, and a concomitant upregulation of fibroblastic markers such as type I collagen, ultimately leading to the formation of mechanically inferior fibrocartilage [3]. Numerous strategies have been explored to mitigate dedifferentiation during chondrocyte expansion, including supplementation with growth factors such as fibroblast growth factors (FGFs), transforming growth factor-β (TGF-β), insulin-like growth factors, and bone morphogenetic proteins (e.g., BMP-2). While these bioactive molecules play important roles in cartilage development and repair, their clinical translation has been limited by inconsistent efficacy and adverse effects, including chondrocyte hypertrophy, cartilage mineralization, fibrosis, and accelerated cellular aging [4–6].

Alternative approaches, such as co-culture systems involving chondrocytes and other cell types, have shown potential in preserving the chondrocytic phenotype; however, these methods are constrained by increased donor-site morbidity and limited cell availability [7,8]. Other strategies aimed at reversing dedifferentiation, including culture on non–tissue culture–treated substrates [9] or transfer into three-dimensional scaffolds such as alginate hydrogels [10], have demonstrated partial success. Nevertheless, these techniques often require extensive cell manipulation, prolonged culture periods, and complex processing steps, which limit scalability and translational applicability. Collectively, these limitations underscore the need for biomaterial-based solutions that can efficiently modulate cell behavior while minimizing exogenous interventions.

The extracellular matrix plays a central role in regulating tissue-specific structure and function by providing both mechanical support and bioactive cues that govern cell adhesion, migration, proliferation, and differentiation [11]. In the context of bioactive materials, increasing attention has been directed toward decellularized cell-secreted matrices (CSM), which preserve the compositional complexity and hierarchical organization of native ECM. CSM derived from various cell sources, including adipose-derived stem cells and bone marrow–derived mesenchymal stem cells, have been shown to support regeneration of multiple tissues, including bone [12,13], skin, cartilage [14], and skeletal muscle [15]. Importantly, these matrices act as instructive microenvironments, presenting endogenous biochemical and biophysical signals that direct cell fate decisions without the need for supraphysiological growth factor supplementation.

Recent studies have demonstrated that ECM derived from primary human chondrocytes retains bioactive cues capable of supporting chondrocyte expansion while reducing dedifferentiation. However, these investigations have primarily focused on phenotypic maintenance and did not comprehensively evaluate the ability of such matrices to promote broader ECM deposition, matrix remodeling, or differentiation-related outcomes [16]. Furthermore, the translational potential of these systems is often limited by challenges related to storage, handling, and reproducibility.

### Study Hypothesis and Objectives

In this study, we hypothesized that a decellularized and lyophilized cell-secreted matrix functions as a bioactive substrate capable of regulating cell behavior by promoting chondrogenic differentiation while attenuating chondrocyte dedifferentiation, without the need for exogenous growth factor supplementation. To test this hypothesis, we generated CSM from rabbit auricular perichondrial (AuP) cells, followed by decellularization and lyophilization to create a stable, storable biomaterial. The resulting matrices were characterized and evaluated for their ability to support chondrocyte expansion, enhance chondrogenic differentiation of progenitor cells, and modulate dedifferentiation processes. Additionally, we assessed the compatibility of this system with allogeneic and xenogeneic cells and introduce a versatile plate-based CSM preparation strategy with broad applicability in bioactive materials and cartilage tissue engineering

## Materials and methods

### Preparation of cell-secreted matrices-coated plates from rabbit auricular perichondrial cells

Rabbit ears were obtained from a 3-week old animals after approval of the procedure by local animal care ethical committee. Perichondrium tissue was dissected from auricular cartilage and cultured as a primary outgrowth for 1 month. Thereafter, the rabbit auricular perichondrial cells (rAuP) were passaged and expanded *in vitro* until passage 3 where they were freezed at -150°C for further use.

For CSM preparation, rAuP cells were seeded at a density of 6.10^5^ per well in a 6-well plates and cultured in chondrogenic medium for 1 month until overconfluency. Chondrogenic medium consisted of DMEM high glucose with glutamine and sodium pyruvate (DMEM, Dutcher, Bernolsheim, France), 0.1% antibiotics, 100nM dexamethasone, 50μg/ml ascorbic acid-2 phosphate, 40 μg/ml proline, 10 ng/ ml of Transforming growth factor beta-3 (TGF beta 3, Sigma-Aldrich, Saint-Quentin-Fallavier, France) and Insulin Transferrin Selenium media supplement, 1X (ITS+1). Cells formed then a cellular layer which was decellularized within the plate by adding 1% sodium dodecyl sulfate (SDS) under agitation for 24 hours. Thereafter, the plates were washed thoroughly with distilled water to remove traces of SDS. Then, the plates containing the acellular layer was lyophilized in the freeze-dryer CHRIST Alpha1/2,4 (Grosseron, Couëron, France) for 1 hour at 1 mbar and stored at room temperature for further use.

### Isolation and cultivation of human cartilage cells

Human cartilage was obtained from the femoral heads of different patients (average age : 55 years) undergoing hip replacement surgery. The experimental protocol was approved by local ethical committee, named comité de protection des personnes Nord Ouest III, in accordance with the applicable guidelines and regulations. All patients provided signed consent agreement before surgery. Cells were released from cartilage digestion using 2 mg/ml of protease type XIV for 30 min (Sigma-Aldrich) and 2 mg/ml of collagenase type II overnight (Thermo Fisher Scientific, Illkirch-Graffenstaden, France). The cells were then washed with PBS, seeded at a density of 4.10^4^ cells/cm^2^ and cultured in Dulbecco’s modified Eagle medium (DMEM, Dutscher), supplemented with 10% fetal bovine serum (FBS, Dutscher), 100 IU/ml of streptomycin (Lonza, Levallois Perret, France), 0.25μg/ml of fungizone (Sigma-Aldrich) and then incubated at 37°C in a humidified atmosphere containing 5% CO2. Media was changed twice a week.

Human AuP were derived from auricular cartilage biopsies after 12-year-old children undergone rhinoplasty. These surgical wastes were obtained at the maxillofacial surgery surgical unit of the Centre Hospitalier Universitaire (CHU) of Caen, from patients (or legals) who signed consent forms. Perichondrium was separated from cartilage by dissection and sliced into small pieces. Cells were obtained by sequential enzymatic digestion using pronase for 45-60 min at 37°C and collagenase II (Thermo Fisher Scientific) overnight at 37°C. The next day, the cells were collected after centrifugation and seeded in T75 flasks and amplified until reaching passage 3, then freezed for further use.

### Stably GFP expressing cells

Stably GFP-transfected rabbit perichondrial cells were obtained by transfection with pEGFP-N1 plasmid (BD Biosciences Clontech, Le Pont-de-Claix, France) and G418 (Sigma-Aldrich) antibiotic selection. Stably transfected human cells were generously given by Pr S. Allouche (CHU, Caen).

### Fluorescence and normal microscopy

Cultured cells were fixed in 10% Neutral Buffered Formalin (NBF, Sigma-Aldrich) at 4°C for 20 min, washed twice with phosphate-buffered saline (PBS, Lonza), and then stained with 4,6-diamidino-2-phenylindole (DAPI, Sigma Aldrich) to identify nuclear components. The stained samples, as well as microphotographs of cells that have been taken at different passages were examined using EVOS cell imaging system M7000 (Thermo Fisher Scientific).

### Immunohistochemistry of type II collagen

After cultures and fixation with NBF, the plates were rinsed in PBS and incubated overnight at 4°C with anti-collagen II primary antibody (1:100, ab 34712, abcam, Cambridge, UK). A negative control was performed by replacing primary antibody solutions with PBS. The next day, the plates were rinsed five times in PBS to remove traces of primary antibody and a secondary antibody (Alexa Fluorophore 594 conjugated anti-rabbit, Thermo Fisher Scientific) diluted at 1:800, was added for 1.5 hour. After rinsing three time with PBS, microphotographies were done on EVOS cell imaging system microscope M7000.

### Histological staining

Histological stainings were performed directly on culture plates. For Masson’s trichrome staining, samples were incubated with Masson’s solution composition for 5 min. Phosphomolybdic acid was next added for 5 min before rinsing with 0.2% acetic acid. The last step consisted on incubating samples with aniline blue for 5 min and excess stain was removed by rinsing with 0.2% acetic acid. For Safranin O staining, samples were incubated in 0.1% Safranin O solution for 5 min. For Hematoxylin Eosin staining, samples were incubated with Harris-modified Hematoxylin solution for 3min and then switched to Eosin solution for 3 min.

Alcian blue was also performed on the plates, by incubation with 3% acetic acid at pH 2.5 for 3min, then 1% Alcian blue at pH 2.5 for 45min before rinsing twice with distilled water for 10 min. The plates were then prepared for macroscopic observation and microphotography under Zeiss Axioscope and ZEN software.

### Protein extraction and biochemical analysis

Total proteins were extracted using radio-immunoprecipitation assay (RIPA) lysis buffer supplemented with a cocktail of phosphatase (Na_2_VO4, 10μg/ml) and protease inhibitors (1μg/ml pepstatin A, 1μg/ml aprotinin, 4μg/ml phenylmethlsulfonyl floride and 1μg/ml leupeptin). Fastin™ Elastin, Sircol ™ soluble collagen and Blyscan™ sulfated Glycosaminoglycans assay kits (Biocolor Life Sciences Assays, Carrickfergus, UK) were used to quantify elastin, collagen and GAGs, respectively, according to the manufacturer’s protocols.

### Proteomic analysis

Protein extracts (5 µg of each) extracted by RIPA buffer in absence of any protease inhibitors were prepared using a modified Gel-aided sample preparation protocol[17]. Samples were digested with trypsin/Lys-C overnight at 37°C. For nano-LC fragmentation, protein or peptide samples were first desalted and concentrated onto a µC18 Omix (Agilent, Les Ulys, France) before analysis.

The chromatography step was performed on a NanoElute (Bruker Daltonics, Wissembourg, France) ultra-high-pressure nano flow chromatography system. Approximatively 200 ng of each peptide sample were concentrated onto a C18 pepmap 100 (5mm x 300µm i.d.) precolumn (Thermo Fisher Scientific) and separated at 50°C onto a reversed phase Reprosil column (25cm x 75μm i.d.) packed with 1.6μm C18 coated porous silica beads (Ionopticks, Australia). Mobile phases consisted of 0.1% formic acid, 99.9% water (v/v) and 0.1% formic acid in 99.9% ACN (v/v). The nanoflow rate was set at 400 nl/min, and the gradient profile was as follows: from 2 to 15% B within 60 min, followed by an increase to 25% B within 30 min and further to 37% within 10 min, followed by a washing step at 95% B and reequilibration.

Mass spectroscopy (MS) experiments were carried out on an TIMS-TOF pro mass spectrometer (Bruker Daltonics) with a modified nano electrospray ion source (CaptiveSpray, Bruker Daltonics). The system was calibrated each week and mass precision was better than 1 ppm. A 1400 spray voltage with a capillary temperature of 180°C was typically employed for ionizing. MS spectra were acquired in the positive mode in the mass range from 100 to 1700 m/z. In the experiments described here, the mass spectrometer was operated in PASEF mode with exclusion of single charged peptides. A number of 10 PASEF MS/MS scans was performed during 1.25 seconds from charge range 2-5.

Before post-process, the samples are analysed using Preview software (ProteinMetrics) in order to estimate the quality of the tryptic digestion and predict the post-translational modifications present. The fragmentation pattern was used to determine the sequence of the peptide. Database searching was performed using the Peaks X+ software. A UniProt *Oryctolagus cuniculus* database was used. The variable modifications allowed were as follows: K-acetylation, methionine oxidation, Deamidation (NQ) and Methylation (KR). In addition, C-Propionoamide was set as fix modification. “Trypsin” was selected as Specific. Mass accuracy was set to 30 ppm and 0.05 Da for MS and MS/MS mode, respectively. Data were filtering according to a FDR of 0.1%, 2 unique peptides and the elimination of protein redundancy on the basis of proteins being evidenced by the same set or a subset of peptides. Enrichment analyses were perfomed with KEGG pathways database and Gene Ontology resource and allowed to describe the molecular functions, biological processes, and primary cellular localization.

### RNA extraction and real time reverse-transcriptase polymerase chain reaction

Total RNA was purified using Qiagen RNeasy® mini kit (Qiagen, Courtaboeuf, France), according to the manufacturer’s protocol. Total RNA (1μg) was treated with DNAse before reverse transcription into cDNA using Moloney Murine Leukemia Virus M-MLV (Invitrogen by Thermo Fisher Scientific) and oligodT primers. Specific transcripts were then amplified by real time PCR on StepOnePlus apparatus (Applied Biosystems by Thermo Fisher Scientific) using Power SYBER Green mix (Thermo Fisher Scientific) and the following human primer sequences. The expression of each gene was normalized to three housekeeping genes (GAPDH, RPL13 and β2Microgmobulin):

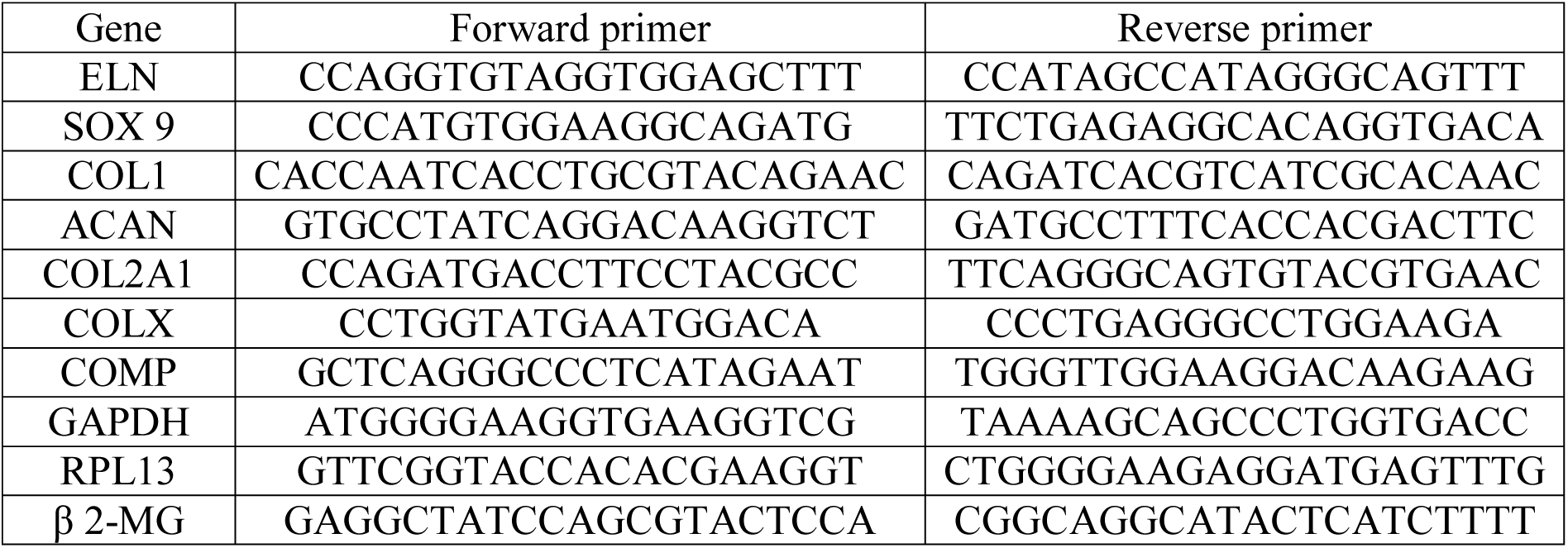

### Cell viability Assay

The colorimetric WST-1 kit (Roche Diagnostics, Meylan, France) was used for viability tests. The amount of formazan formed is directly related to the metabolic activity of cells. Chondrocytes were seeded on 6-well plates (either plastic or CSM coated plates). After several days of culture, WST-1 reagent was added for 3 hours. Absorbance was measured on aliquot fractions of the supernatants at 450 and 650 nm. The same procedure was repeated after 1 (P1) or 3 (P3) passages. The results were normalized and expressed as relative to controls that were measured in plastic plates (control).

### Statistical analyses

Data were presented as mean ± standard deviation (S.D.). Differences between groups was observed using two-way analysis of variance with Tukeys’s multiple comparison tests using Prism software. Other sample groups were subjected to unpaired t-test. Values less than 0.05 were considered as significant. *: P≤0.05; **: P≤0.01; ***P≤0.001.

## Results

### Generation and Structural Characterization of Cell-Secreted Matrices

Cell-secreted matrices (CSM) were generated by culturing rabbit auricular perichondrial (rAuP) cells beyond confluency to promote extracellular matrix deposition. The resulting cell layers were decellularized using sodium dodecyl sulfate (SDS) aiming of removing cellular components while preserving matrix architecture. Successful decellularization was confirmed by the complete absence of nuclei following DAPI staining (Fig. 1A). Subsequent lyophilization produced a uniform, whitish coating adherent to the culture plate surface, indicating preservation of matrix integrity after freeze-drying (Fig. 1B).

**Figure 1.**
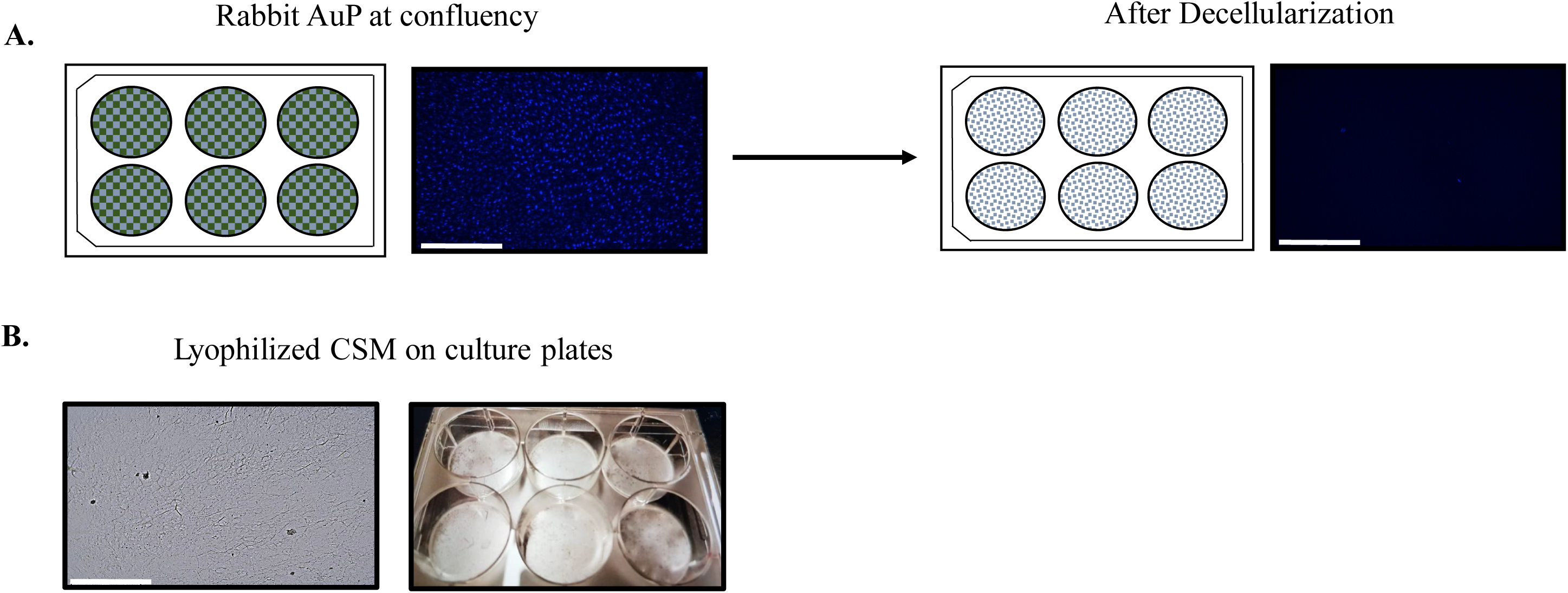
Preparation of rabbit cell-secreted matrices (CSM) A. DAPI staining before and after decellularization of cell layers of confluent rabbit auricular perichondrial cells (AuP). B. Microscopic and macroscopic aspects of lyophilized CSM showing the remaining secreted matrices as white coating. Scale bar: 500μm.

Histological and cytochemical analyses further confirmed the effective removal of cellular material and retention of ECM components. No nuclear structures were detected across multiple stainings (Fig. 2), supporting the robustness of the decellularization process. Masson’s trichrome staining demonstrated strong preservation of collagen fibers, as evidenced by sustained dark blue staining following both decellularization (Fig. 2b) and lyophilization (Fig. 2c). In contrast, sulfated glycosaminoglycans (GAGs), visualized by Safranin O staining, were partially reduced after decellularization and further diminished following lyophilization (Fig. 2e,f), indicating greater sensitivity of proteoglycan-associated components to processing. Hematoxylin and eosin staining revealed retention of extracellular proteic material, although staining intensity was reduced relative to native layers, likely reflecting the loss of cell-associated matrix components (Fig. 2h,i).

**Figure 2.**
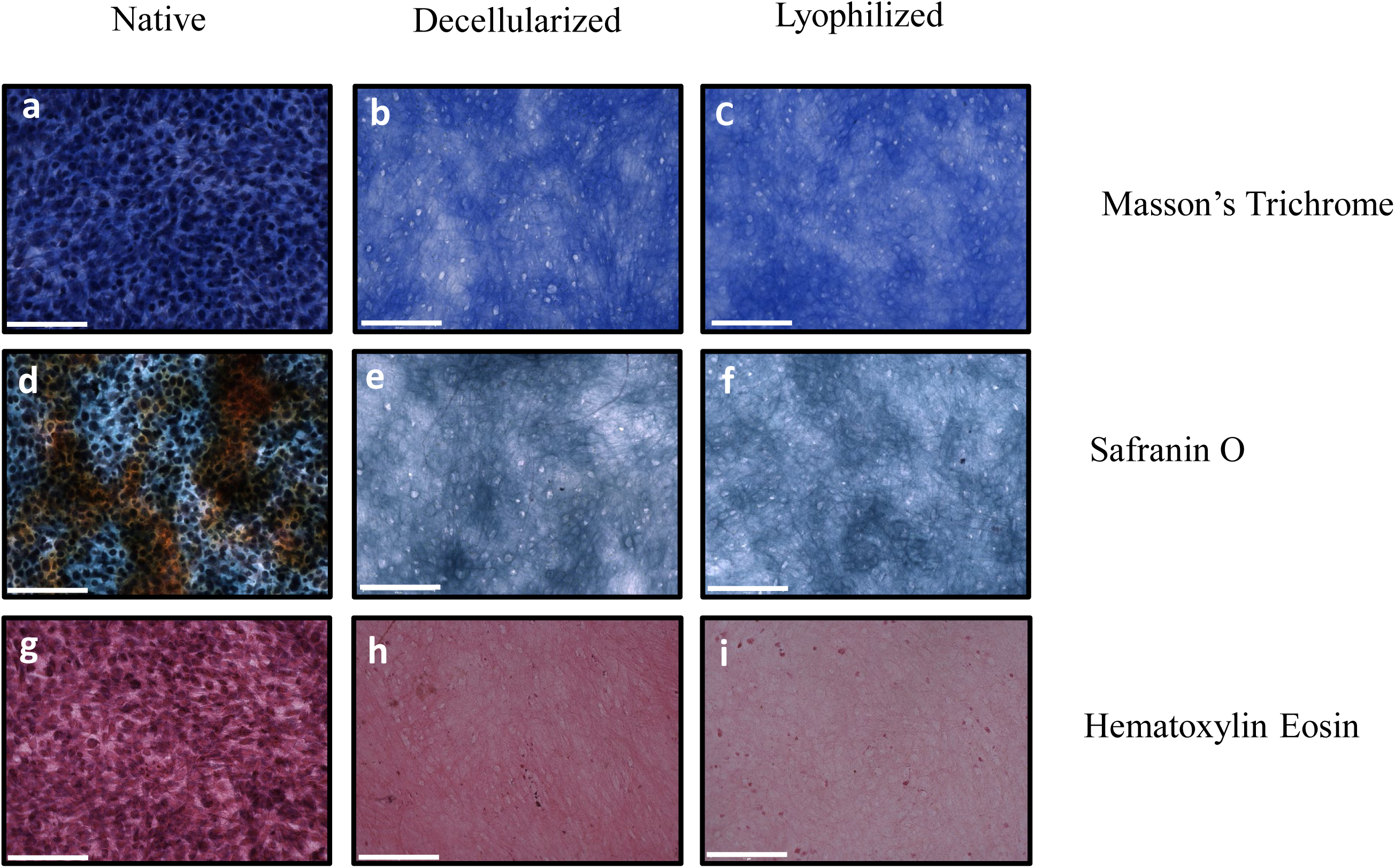
Histological characterization of CSM before and after decellularization and lyophilization. Histological staining of collagen fibers using Masson’s Trichrome (**a**,**b**,**c**), glycosaminoglycans by Safranin O staining (**d,e,f**) and extracellular proteins using Hematoxylin Eosin (**g,h,i**). Scale bar: 200μm.

### Quantitative Protein Composition and Proteomic Profiling of Lyophilized CSM

To quantify matrix retention, total protein content was measured and normalized to surface area. Native AuP cell layers contained approximately 1.5 µg/cm² of total protein, which decreased by approximately 50% following decellularization and further declined to 0.6 µg/cm² after lyophilization (Fig. 3). Targeted biochemical assays demonstrated similar proportional reductions in elastin and GAG content, whereas total collagen content exhibited a comparatively smaller decrease, indicating preferential preservation of fibrillar collagen during the process.

**Figure 3.**
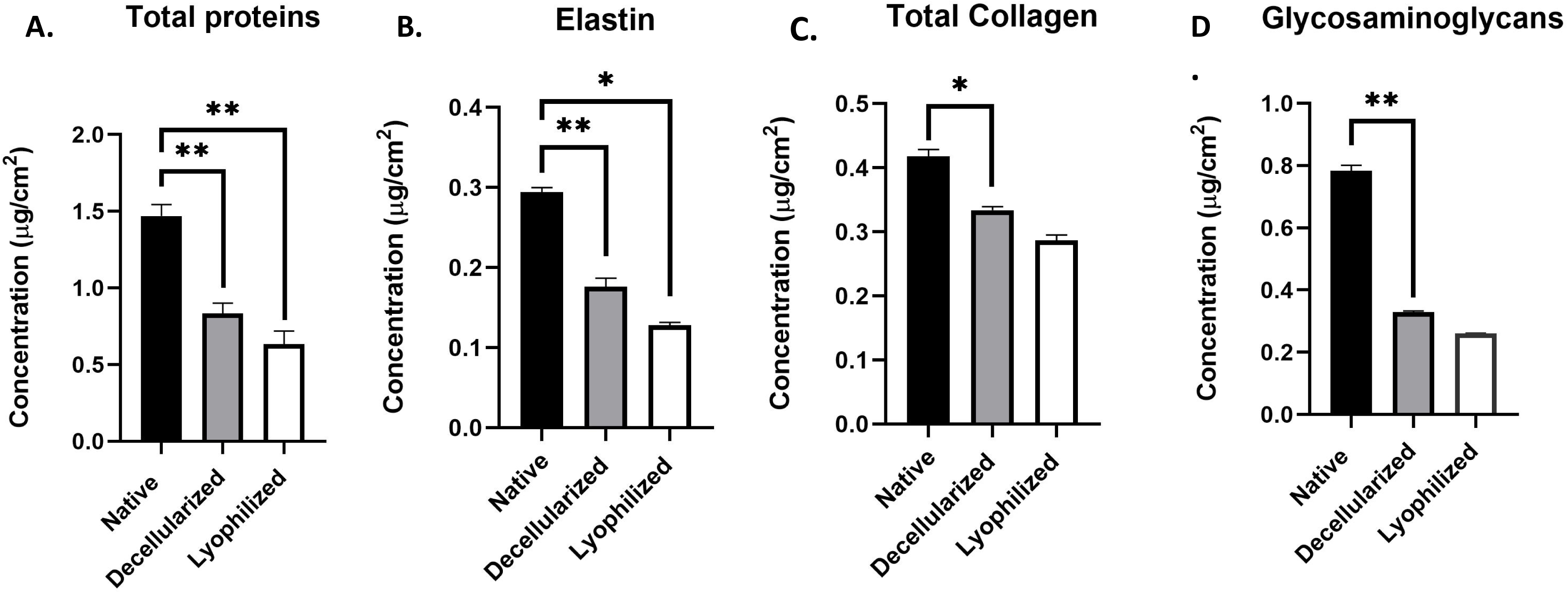
Biochemical characterization of CSM. Protein concentration was determined by Bradford assay (A), ELN content by Fastin assay (B), Total Collagen was measured by Sircol assay (C), and sulfated GAG’s by Blyscan assay (D).

To assess matrix composition and batch-to-batch reproducibility, proteomic analysis was performed on six independently prepared lyophilized CSM samples. Mass spectrometry followed by stringent filtering identified a core set of 46 proteins consistently present across all samples. Proteins were normalized to molecular weight and ranked by relative abundance (Table 1). Gene Ontology (GO) and KEGG pathway enrichment analyses revealed extracellular matrix organization as the most significantly enriched biological process. Structural ECM proteins—including fibrillin-1, fibronectin, collagens I and XII, biglycan, decorin, dermatopontin, and tenascin-C—dominated the proteomic profile, driving strong enrichment of extracellular matrix and extracellular region cellular component terms.

**Table 1:** Set of 46 common proteins found in all samples by proteomics and sorted by their relative abundance.

| Protein number | Accession | Description | Score | Primary localization |
| --- | --- | --- | --- | --- |
| 1 | G1SKM2 | Fibrillin 1 | 238,45 | Extracellular matrix (ECM) |
| 2 | G1SWS9 | Vimentin | 235,53 | Cytoskeleton (cytoplasmic intermediate filaments) |
| 3 | A0A5F9CJW0 | Fibronectin | 190,5 | Extracellular matrix (ECM) |
| 4 | G1U9I8 | Cytokeratin-1 | 180,28 | Cytoskeleton (keratin intermediate filaments) |
| 5 | G1T1V0 | Keratin 10 | 179,87 | Cytoskeleton (keratin intermediate filaments) |
| 6 | A0A5F9D471 | 2-phospho-D-glycerate hydro-lyase | 179,45 | Cytosol |
| 7 | A0A5F9DEV8 | Annexin | 176,35 | Cytosol / plasma membrane-associated |
| 8 | G1U4C2 | Coiled-coil domain containing 80 | 173,26 | Extracellular matrix / secreted |
| 9 | G1SH05 | Tubulin beta chain | 164,63 | Cytoskeleton (microtubules) |
| 10 | G1SRW4 | Elastin microfibril interfacer 1 | 159,09 | Extracellular matrix (ECM) |
| 11 | P68105 | Elongation factor 1-alpha 1 | 157,59 | Cytosol |
| 12 | A0A5F9CJB7 | Collagen alpha-1(XII) chain | 157,03 | Extracellular matrix (ECM) |
| 13 | A0A5F9DFJ9 | Glyceraldehyde-3-phosphate dehydrogenase | 150,65 | Cytosol |
| 14 | G1TPZ1 | Galectin | 147,39 | Cytosol / extracellular |
| 15 | A0A5F9CGP0 | Histone H4 | 134,22 | Nucleus (chromatin) |
| 16 | G1SXR1 | Proline and arginine rich end leucine rich repeat protein | 132,9 | Nucleus / cytosol |
| 17 | A0A5F9DUM8 | Heat shock protein family A (Hsp70) member 8 | 129,6 | Cytosol (also nucleus) |
| 18 | U3KML1 | Biglycan | 128,56 | Extracellular matrix (ECM) |
| 19 | G1SNE8 | Calpactin I light chain | 123,95 | Cytosol / plasma membrane-associated |
| 20 | Q9TTC6 | Peptidyl-prolyl<br>cis-trans isomerase A | 123,67 | Cytosol |
| 21 | G1TKL2 | Uncharacterized<br>protein | 122,03 | Unknown<br>(likely cytosolic) |
| 22 | G1T5H0 | HtrA serine<br>peptidase 1 | 121,56 | Extracellular<br>/ secreted |
| 23 | A0A5F9D8N0 | Histone H2A | 120,9 | Nucleus<br>(chromatin) |
| 24 | A0A5F9DQH9 | Latent<br>transforming growth<br>factor beta binding<br>protein 1 | 117,33 | Extracellular<br>matrix (ECM) |
| 25 | A0A5F9C804 | 78 kDa glucose-<br>regulated protein | 115,58 | Endoplasmic<br>reticulum |
| 26 | G1SER8 | Profilin | 110,65 | Cytosol<br>(actin cytoskeleton) |
| 27 | G1SVK5 | Protein S100 | 107,81 | Cytosol /<br>nucleus |
| 28 | G1SDA2 | Cellular retinoic<br>acid-binding protein 1 | 106,43 | Cytosol /<br>nucleus |
| 29 | P51662 | Annexin A1 | 105,88 | Cytosol /<br>plasma membrane |
| 30 | A0A5F9DPN4 | Tenascin C | 105,73 | Extracellular<br>matrix (ECM) |
| 31 | A0A5F9CXS8 | Collagen alpha-1(I) chain | 100,85 | Extracellular<br>matrix (ECM) |
| 32 | G1T7Z6 | Phosphoglycerate<br>kinase | 100,01 | Cytosol |
| 33 | A0A5F9CGT7 | Heat shock<br>protein HSP 90-beta | 99,63 | Cytosol |
| 34 | G1U155 | Histone H2B | 99,53 | Nucleus<br>(chromatin) |
| 35 | G1SYD6 | Lamin A/C | 95,37 | Nucleus<br>(nuclear lamina) |
| 36 | A0A5F9D1L2 | Transforming<br>growth factor-beta-<br>induced protein ig-h3 | 94,01 | Extracellular<br>matrix (ECM) |
| 37 | P49065 | Albumin | 90,24 | Extracellular<br>/ blood plasma |
| 38 | P01948 | Hemoglobin<br>subunit alpha-1/2 | 88,73 | Cytosol<br>(erythrocytes) |
| 39 | P62160 | Calmodulin | 84,91 | Cytosol /<br>nucleus |
| 40 | G1T2C4 | Transgelin | 77 | Cytoskeleton<br>(actin-associated) |
| 41 | Q28888 | Decorin | 76,37 | Extracellular<br>matrix (ECM) |
| 42 | P18287 | Apolipoprotein E | 75,17 | Extracellular<br>/ plasma /<br>lipoproteins |
| 43 | P29562 | Eukaryotic<br>initiation factor 4A-I<br>(Fragment) | 73,71 | Cytosol |
| 44 | G1SNC7 | Dermatopontin | 73,12 | Extracellular<br>matrix (ECM) |
| 45 | U3KNZ4 | Nucleoside diphosphate kinase | 70,34 | Cytosol |
| 46 | P02099 | Hemoglobin subunit gamma | 62 | Cytosol |

Processes related to collagen fibril organization, cell–matrix adhesion, and regulation of transforming growth factor–β signaling were also significantly overrepresented, consistent with the presence of matrix-associated regulatory proteins such as latent TGF-β–binding protein 1 and TGF-β–induced protein ig-h3. Cytoskeletal organization and actin dynamics pathways were moderately enriched, driven by proteins including vimentin, tubulin, keratins, profilin, and transgelin, while glycolytic enzymes and molecular chaperones accounted for minor contributions. Nuclear and serum-derived proteins were detected at low abundance and contributed minimally to enrichment signals. Collectively, these findings indicate that the lyophilized CSM is highly enriched in structurally and functionally relevant ECM components with limited intracellular contamination.

### CSM Supports Allogeneic and Xenogeneic Cell Adhesion and Expansion

To evaluate the biological compatibility of rabbit-derived CSM, both rabbit allogeneic and human xenogeneic GFP-expressing cells were seeded onto CSM-coated plates. Fluorescence imaging over several days revealed robust cell adhesion, spreading, and morphological adaptation for both cell types (Fig. 4). These results demonstrate that rabbit CSM provides a permissive and supportive substrate for both allogeneic and xenogeneic cell culture applications.

**Figure 4.**
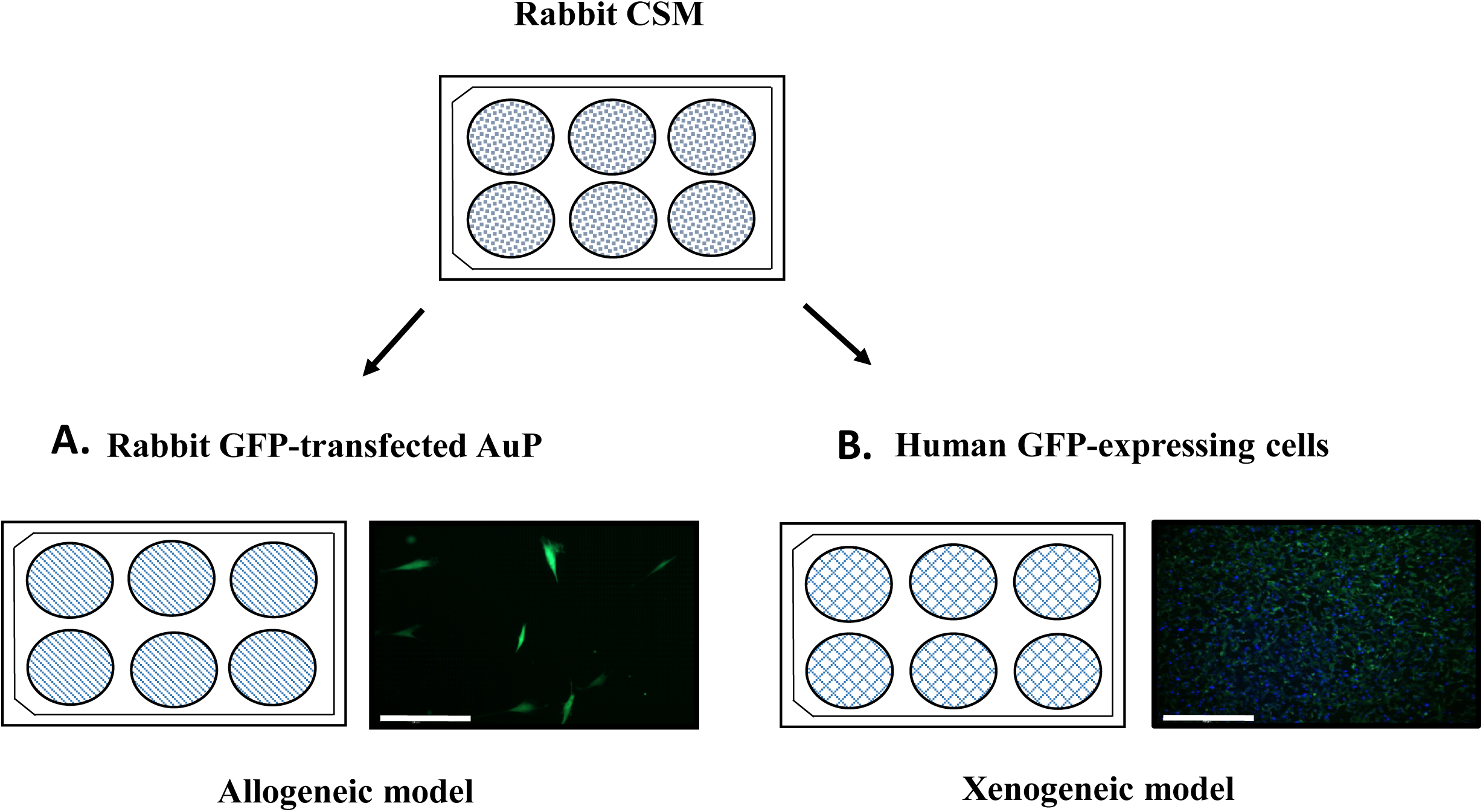
Rabbit CSM as a support for allogenic and xenogeneic culture cells. rAuP cells were cultured until confluency, before decellularization and lyophilization. Cell-Secreted Matrices were then seeded for cell culture and expansion with rabbit (A) and human (B) GFP expressing cells. Fluorescent imaging for DAPI and GFP was performed after 48h of culture.

### CSM Enhances Proliferation of Human Articular Chondrocytes

The impact of CSM on chondrocyte expansion was assessed by subculturing human articular chondrocytes (HAC) on either standard tissue culture plastic or CSM-coated plates from primary culture (P0) through passage 5 (P5). Phase-contrast microscopy revealed consistently higher cell density and more rapid proliferation on CSM-coated surfaces at all passages (Fig. 5A). These observations were corroborated by DAPI nuclear staining up to passage 3 (Fig. 5B). Quantitative assessment using the WST-1 metabolic assay confirmed significantly enhanced cell viability and proliferation on CSM relative to plastic controls (Fig. 5C).

**Figure 5.**
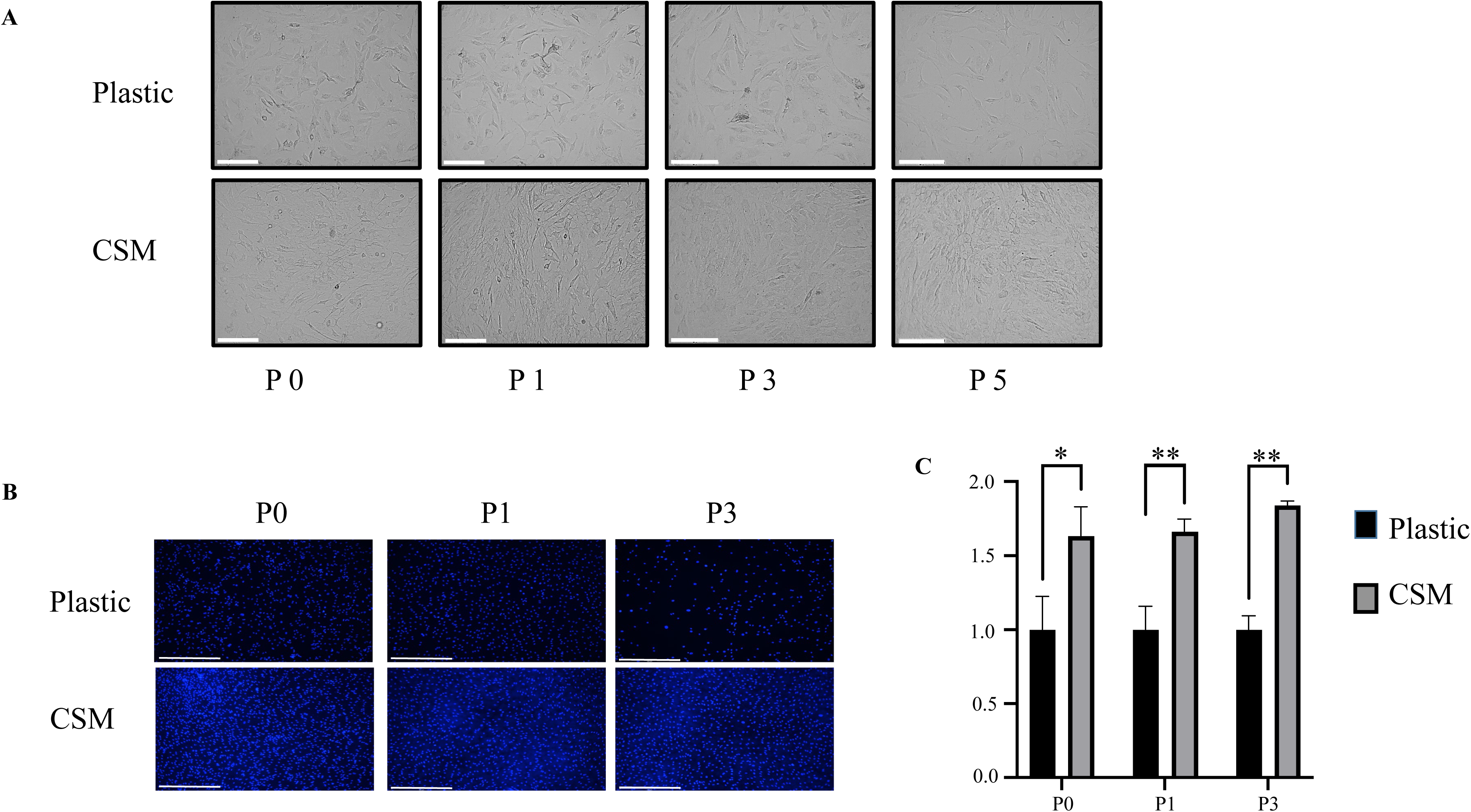
Subculture on CSM promotes cell proliferation. Human articular chondrocytes were subcultured either on plastic and CSM plates from passage 0 until passage 5. **A** : Macroscopic images showing chondrocytes morphology at different passages. **B :** At passages 0, 1 and 3, the cells were fixed and stained with DAPI. **C** : Viability of chondrocytes cultured on CSM plates was measured after passage P0, P1 and P3 via WST-1 test and normalized to control cultures on plastic. (*p˂0.05, **p˂0.01)

### CSM Promotes Chondrogenic Differentiation of Human Progenitor Cells

The ability of CSM to modulate lineage commitment was evaluated using human auricular perichondrial progenitor cells subjected to chondrogenic induction. Cells cultured on CSM-coated plates exhibited markedly increased glycosaminoglycan deposition, as visualized by Alcian blue staining after one and two weeks, compared to plastic controls (Fig. 6A). Gene expression analysis performed after two weeks revealed significant upregulation of cartilage-associated markers, including COL2A1, ELN, COMP, and SOX9, alongside pronounced downregulation of dedifferentiation- and hypertrophy-associated genes COL1 and COLX (Fig. 6B).

**Figure 6.**
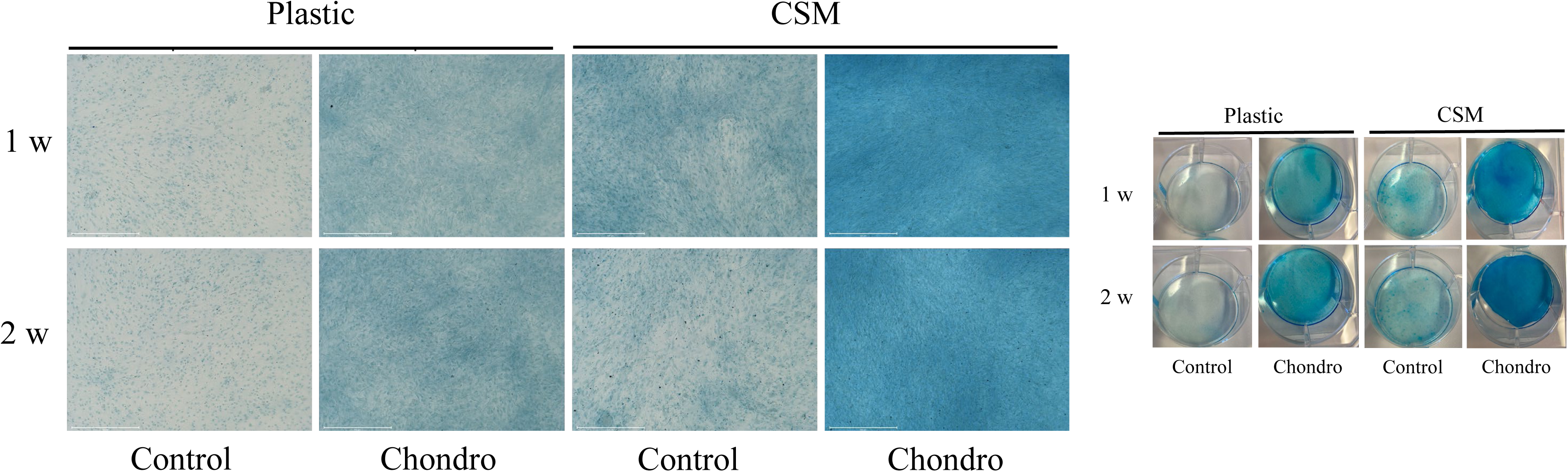
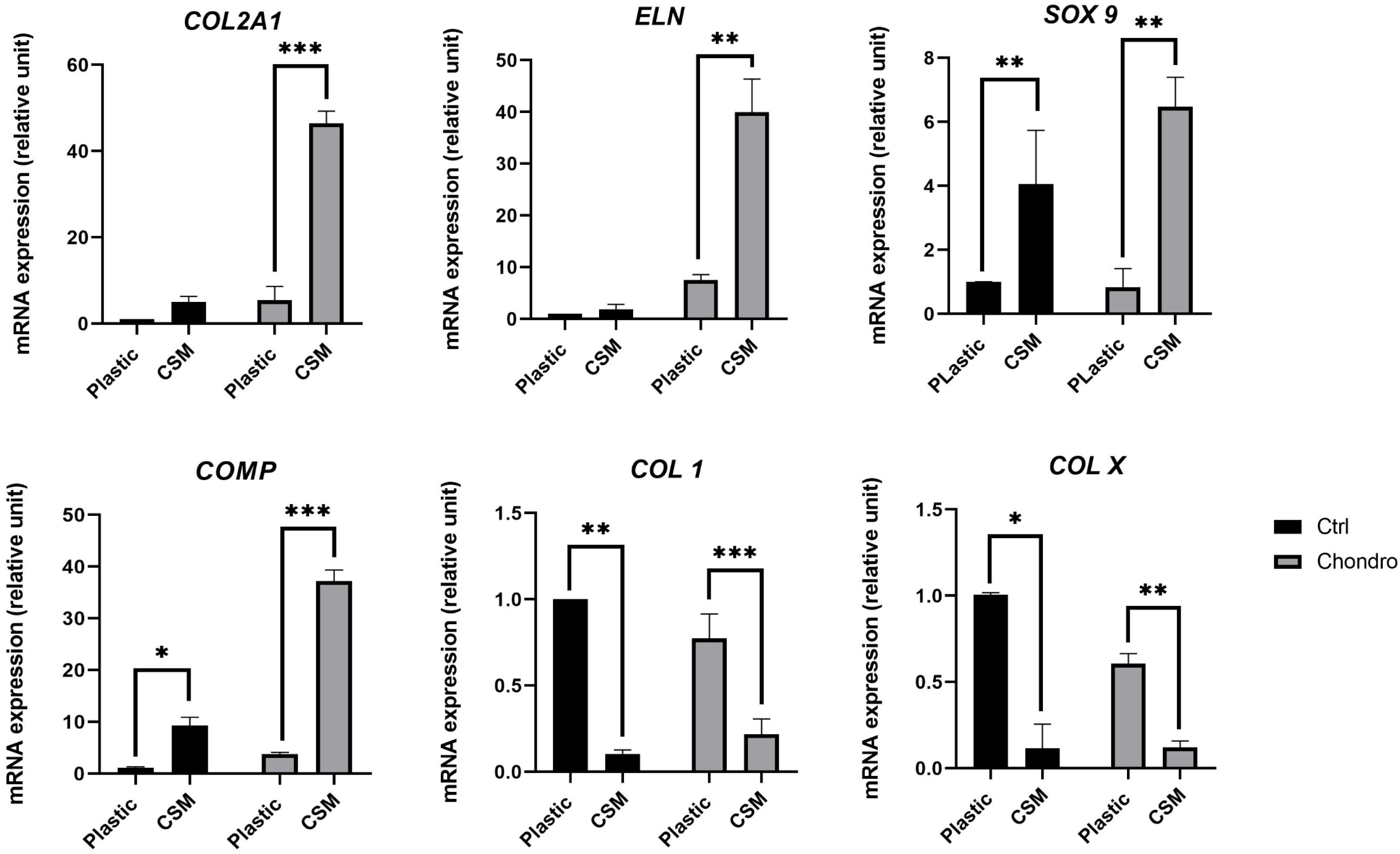
Cell culture on CSM enhances GAG production and chondrogenic differentiation of progenitors cells. Human progenitors from auricular perichondrium were seeded on either plastic or lyophilized CSM plates. At confluency, they were subjected to chondrogenic differentiation. **A :** Alcian Blue staining was done to enlighten GAG production after 1 and 2 weeks. **B** : Gene expression analysis of chondrogenic relevant genes was performed by real-time PCR after 2 weeks of culture. n=3 (*p˂0.05, **P˂0.01, ***p˂0.005).

Notably, enhanced chondrogenic marker expression was observed even under control (non-chondrogenic) culture conditions, indicating that the lyophilized CSM intrinsically provides sufficient biochemical cues to initiate chondrogenic differentiation. These effects were further amplified upon exposure to chondrogenic medium, suggesting a synergistic interaction between matrix-derived signals and soluble factors.

### CSM Preserves the Chondrocytic Phenotype during Serial Expansion

To determine whether CSM could stabilize chondrocyte phenotype during prolonged expansion, primary HAC from three independent donors were serially passaged on either CSM-coated or plastic surfaces through passage 5 (Fig. 7). Cells expanded exclusively on CSM displayed significantly elevated expression of chondrogenic genes ACAN, COL2A1, and SOX9, while expression of the fibroblastic marker COL1 remained comparatively low. These effects became increasingly pronounced with successive passages, with COL2A1 expression at passage 5 exceeding that observed at primary culture by approximately 30-fold.

**Figure 7.**
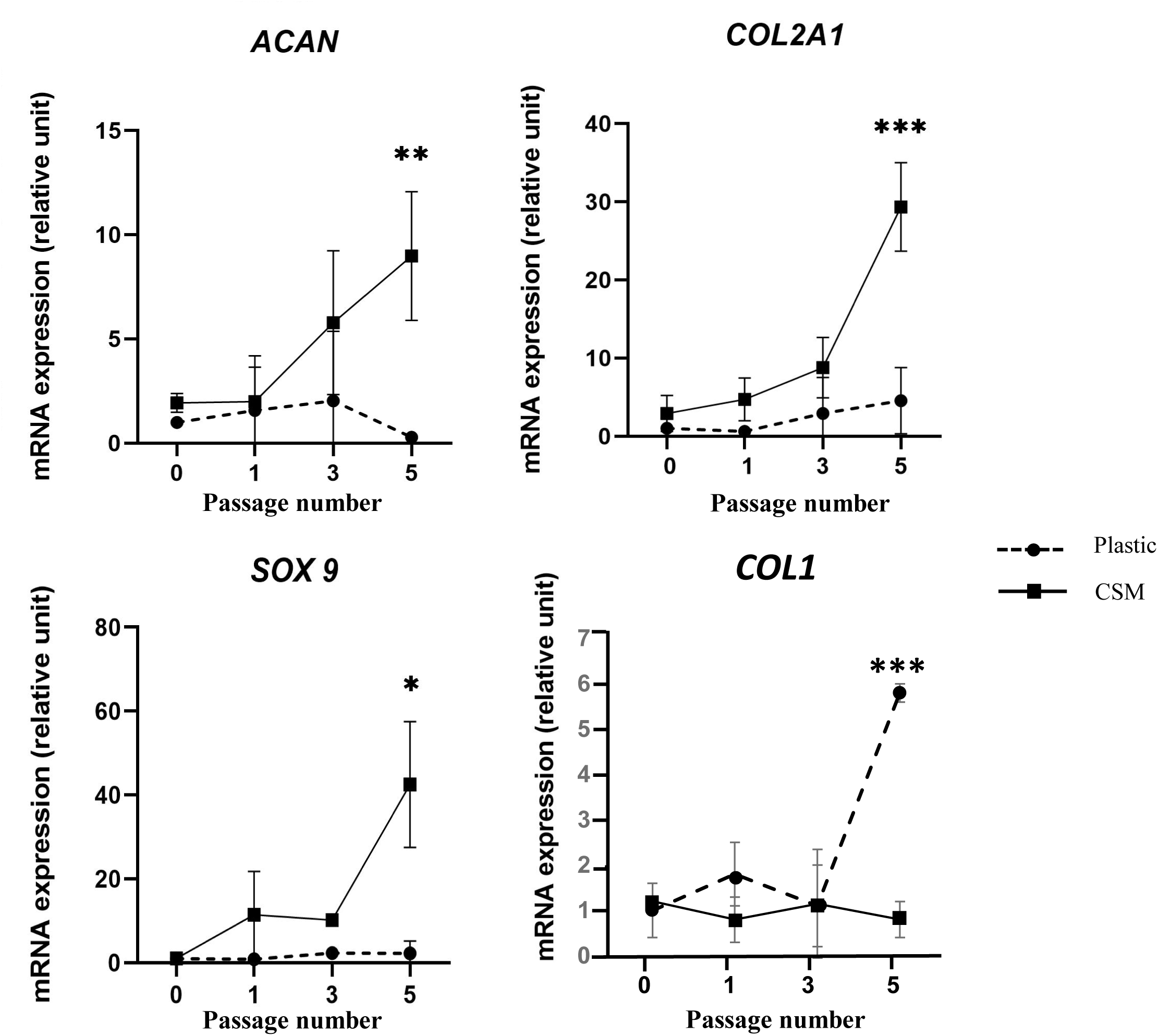
Subculture on CSM maintains chondrogenic phenotype. Human articular chondrocytes were subcultured either on plastic and CSM plates from passage 0 until passage 5. At each passage, cartilage gene markers expression was analyzed by real time RT-PCR, (n=3, *p˂0.05, **p˂0.01, ***p˂0.005).

Immunocytochemical analysis of type II collagen confirmed these transcriptional findings at the protein level. While chondrocytes cultured on plastic exhibited reduced collagen II staining at passage 5, cells maintained on CSM retained strong and widespread collagen II expression (Fig. 8), consistent with sustained chondrocytic identity.

**Figure 8.**
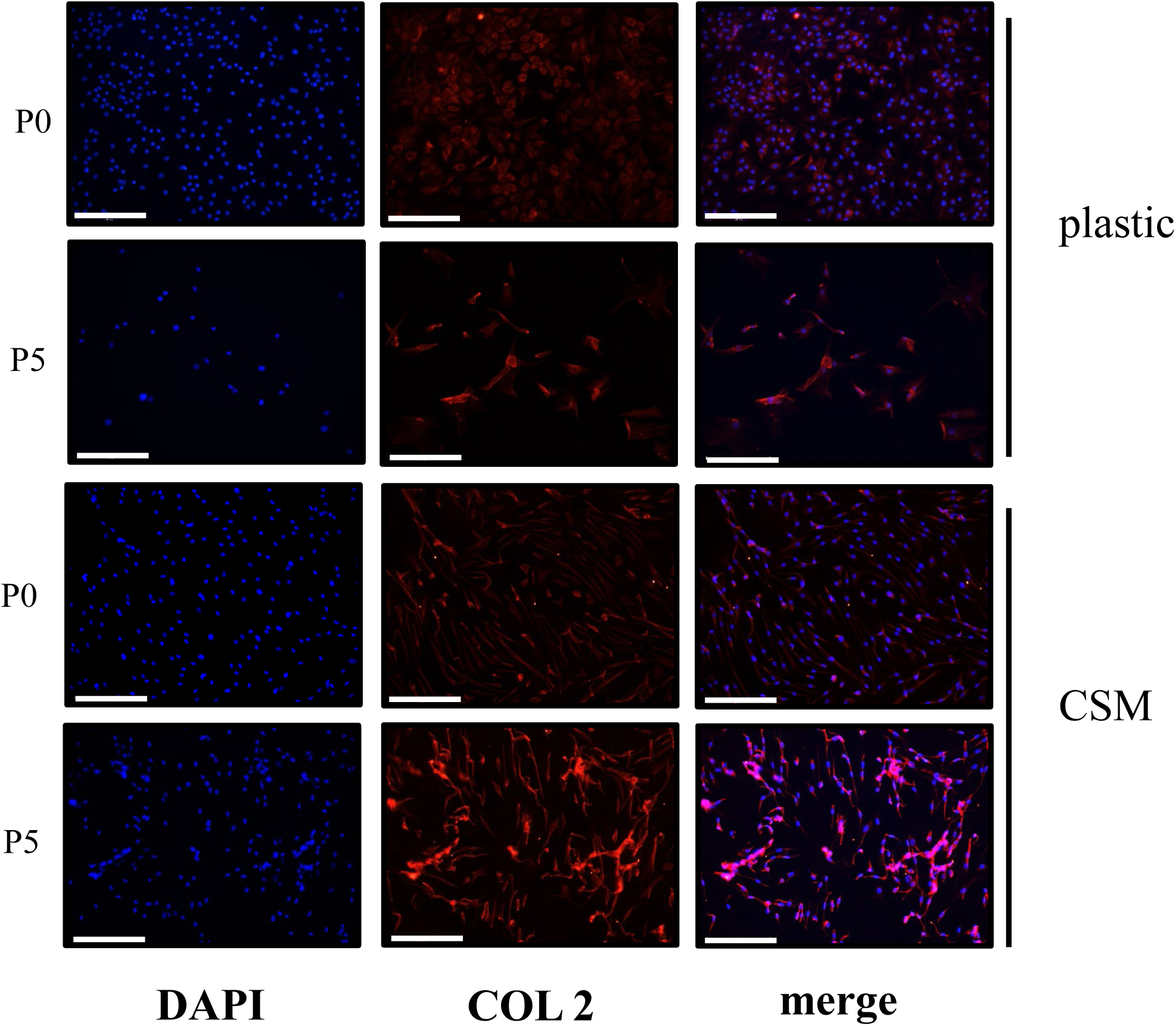
Immunodetection of type 2 collagen upon passages on CSM Vs plastic plates. Comparison of type 2 collagen expression by articular chondrocytes cultured on CSM vs plastic revealed by immunochemistry at passage 0 and passage 5. Scale bar: 200μm.

### CSM Reverses Chondrocyte Dedifferentiation

Finally, we investigated whether CSM could reverse established chondrocyte dedifferentiation. HAC were first expanded on plastic for three passages to induce dedifferentiation, then reseeded onto either plastic or CSM-coated plates for an additional two weeks. Switching dedifferentiated chondrocytes onto CSM resulted in significant re-expression of cartilage-specific genes, including ACAN, COL2A1, and COMP, accompanied by a significative reduction in COL1 and COLX expression (Fig. 9). These findings indicate that CSM not only preserves chondrocyte phenotype during expansion but also partially reverses dedifferentiation, highlighting its potential as a bioactive substrate for cartilage tissue engineering.

**Figure 9.**
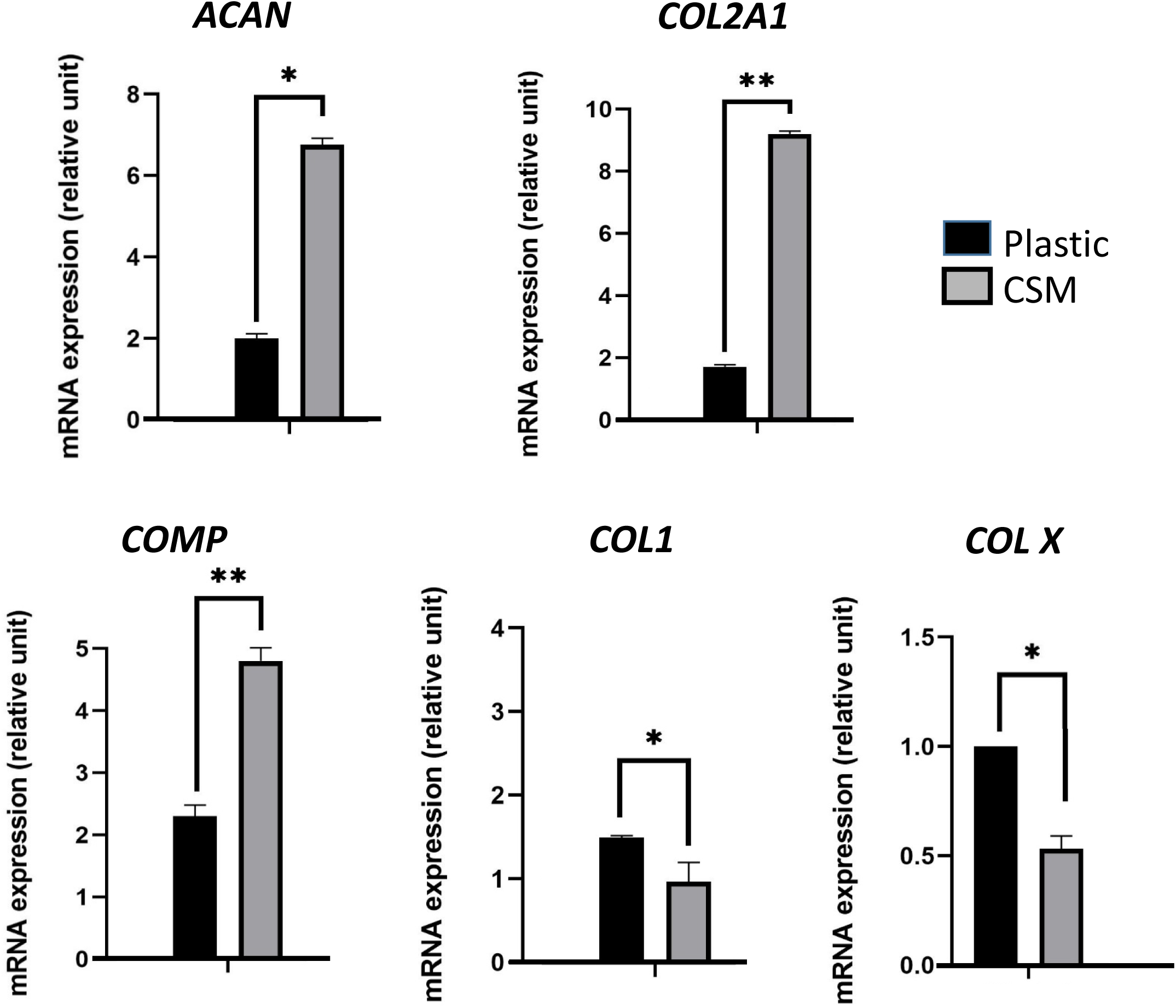
Subculture on CSM reverses the dedifferentiation process. Human articular chondrocytes were subcultured on plastic plates until passage 3. Thereafter, they were trypsinized and seeded either on plastic or switched to CSM plates for 2 additional weeks of culture. Cartilage relevant gene markers expression was then analyzed by real time RT-PCR. (*p˂0.05, **p˂0.01)

## Discussion

The present study demonstrates that a decellularized and lyophilized cell-secreted matrix (CSM) functions as a bioactive material capable of directing cell behavior through intrinsic extracellular matrix (ECM)–derived cues. By preserving a structurally and functionally integrated ECM enriched in fibrillar collagens, proteoglycans, and matricellular proteins, the CSM provides a biologically instructive microenvironment that supports cell adhesion, proliferation, chondrogenic differentiation, and long-term phenotypic stability without reliance on exogenous growth factor supplementation.

Proteomic analysis revealed that the lyophilized CSM is dominated by structural ECM proteins, including fibrillin-1, fibronectin, collagens I and XII, biglycan, decorin, dermatopontin, and tenascin-C. These proteins are well-established regulators of matrix architecture, cell–matrix adhesion, mechanotransduction, and growth factor bioavailability [18–20]. The strong enrichment of extracellular matrix organization and cell–matrix adhesion pathways indicates that the decellularization and lyophilization processes preserved key functional elements of the native matrix while minimizing intracellular debris. Importantly, the presence of regulatory ECM proteins such as latent TGF-β–binding protein 1 (LTBP1) and TGF-β–induced protein ig-h3 suggests that the CSM can locally modulate signaling pathways central to chondrogenesis, aligning with recent findings that ECM sequestration of morphogens enhances regenerative outcomes [21–23].

The enhanced adhesion and proliferation of both allogeneic and xenogeneic cells on rabbit-derived CSM highlight the conserved nature of ECM-mediated signaling. Integrin engagement with matrix ligands such as fibronectin and collagens stimulates focal adhesion assembly and downstream mechanosensitive pathways (FAK, MAPK), which are known to support cell survival and proliferation [24,25]. These interactions likely contribute to the increased metabolic activity observed on CSM-coated surfaces.

A central finding of this work is the ability of CSM to promote chondrogenic differentiation of progenitor cells and maintain the phenotype of mature chondrocytes during extensive expansion. Loss of chondrocyte phenotype during in vitro culture has been attributed to altered mechanotransductive signaling and absence of native pericellular cues when cultured on rigid plastic substrates [26–28]. The observed upregulation of cartilage-specific markers (COL2A1, ACAN, COMP, SOX9) and concurrent downregulation of dedifferentiation markers (COL1, COLX) indicate that CSM restores a tissue-mimicking microenvironment capable of promoting an anabolic cartilage profile, in agreement with recent studies demonstrating that decellularized cartilage matrices support chondrogenesis without exogenous growth factors [29–31].

Notably, chondrogenic responses were observed even in the absence of induction medium, suggesting that CSM provides sufficient biochemical and biophysical cues to initiate lineage commitment. This supports the growing paradigm in bioactive materials research that harnessing matrix-derived signals (e.g., integrin ligands, matricellular domains, bound morphogens) can obviate high-dose recombinant factors, reduce cost, and enhance physiological relevance [32–37].

The ability of CSM to partially reverse established chondrocyte dedifferentiation is particularly significant. Dedifferentiation is associated with disrupted integrin signaling, actin cytoskeleton reorganization, and dysregulated mechanotransductive pathways such as YAP/TAZ and Wnt/β-catenin [38,39]. By reintroducing a biologically relevant ECM environment, CSM appears to reprogram dedifferentiated chondrocytes toward a cartilage-specific transcriptional state. Importantly, the absence of increased COLX expression suggests that redifferentiation occurs without promoting hypertrophy, addressing a key barrier in cartilage tissue engineering.

Although the proteomic analysis detected cytoskeletal proteins and low levels of histones, their presence likely reflects the inherent biological complexity of cell-derived matrices rather than incomplete decellularization. At first glance, the presence of these proteins could be interpreted as incomplete decellularization; however, several lines of evidence argue against this interpretation and support a biologically meaningful origin. First, the absence of detectable nuclei by DAPI staining and the lack of significant enrichment of nuclear biological processes in GO analyses indicate that cellular remnants are minimal.

Cytoskeletal proteins are increasingly recognized as integral components of cell-derived matrices, where they become incorporated into the pericellular and interstitial ECM during matrix deposition, cytoskeletal remodeling, and cell–matrix crosstalk. Vimentin and keratins, in particular, have been shown to associate with focal adhesions and integrin complexes and can persist in decellularized matrices as structural or signaling intermediates. Their retention may therefore contribute to the mechanotransductive properties of the CSM, facilitating cytoskeleton–ECM coupling in newly seeded cells [40,41].

Likewise, histones have been identified in our decellularized ECM preparations. Extracellular or matrix-associated histones have been reported in multiple decellularized matrix systems and are increasingly appreciated for their noncanonical roles. Indeed, beyond their nuclear functions, histones can localize outside the cell and bind ECM components, modulate matrix stiffness, and influence cell adhesion and immune responses [42]. Most of them exist as nucleosomes, as components of extracellular chromatin, as they localize to extracellular vesicles (EVs) and are secreted via the multivesicular body/exosome pathway [43]. However, the lack of nuclear enrichment in GO analysis and absence of nuclear structures by imaging support the conclusion that these components do not represent significant cellular residue.

Taken together, the presence of cytoskeletal and nuclear proteins likely reflects the biological origin and complexity of the cell-secreted matrix rather than a technical limitation of the decellularization process. Their retention may in fact enhance the bioactivity of the material by providing additional structural and signaling cues that are absent in simplified ECM formulations.

Despite the promising findings presented here, several limitations should be acknowledged. First, the proteomic analysis provides a comprehensive qualitative and semi-quantitative overview of CSM composition but does not resolve spatial organization or post-translational modifications of matrix proteins, which are known to critically influence cell behavior. Second, although rabbit-derived CSM supported both allogeneic and xenogeneic cell culture in vitro, in vivo immunological compatibility and long-term host responses were not evaluated and warrant further investigation. Additionally, while the study demonstrates robust effects on chondrocyte phenotype and progenitor differentiation in two-dimensional culture, the performance of CSM in three-dimensional systems or under dynamic mechanical loading conditions remains to be explored. Such conditions more closely recapitulate the native cartilage environment and are highly relevant for translational cartilage repair strategies. Finally, although dedifferentiation reversal was clearly observed at the transcriptional and protein levels, deeper mechanistic interrogation of the signaling pathways involved—such as integrin-mediated mechanotransduction, TGF-β sequestration and release, or YAP/TAZ signaling—was beyond the scope of the current study.

From a translational perspective, the findings of this study open several promising avenues for future research and application. Lyophilization of CSM directly on the plates, represents a significant advantage. Freeze-drying enables long-term storage, ease of transport, and reproducible application while preserving biological activity, consistent with current work on decellularized ECM biomaterials. The plate-based CSM format further simplifies integration into cell manufacturing and assay workflows, mitigating challenges associated with three-dimensional scaffolds and complex growth factor systems. From a biomaterials standpoint, the demonstrated stability and bioactivity of lyophilized CSM position it as a versatile, off-the-shelf matrix coating that could be used in disease modeling platforms, and regenerative medicine workflows.

Future studies could explore scaling and standardization of CSM production, including matrix thickness control, compositional tuning through cell source selection, and batch-to-batch quality metrics aligned with regulatory expectations.

In summary, this study establishes lyophilized CSM as a biologically active material that harnesses native ECM complexity to regulate chondrocyte fate. By promoting chondrogenic differentiation, preserving phenotype during expansion, and reversing dedifferentiation, CSM represents a promising strategy for cartilage tissue engineering. More broadly, these findings reinforce the concept that ECM-derived materials constitute a distinct class of bioactive materials capable of directing cell behavior through intrinsic biological signaling, providing a foundation for next-generation bioactive materials that leverage endogenous signaling rather than exogenous biochemical manipulation.

## Acknowledgement

MH and BD were recipients of PhD grant from University of Caen and Region Normandy, respectively.

## Data statement

The data will be available within a reasonable time after publication, from the corresponding author.

## Declaration of generative AI and AI-assisted technologies in the writing process

During the preparation of this work the authors used OpenAI ChatGPT service in order to improve the readability and language of the manuscript. After using this tool, the authors reviewed and edited the content as needed and take full responsibility for the content of the published article.

## Author contributions

**MH :** Investigation, Original draft, Formal analysis, **BD** : Investigation, **AV** : Investigation, resources, **BB** : Investigation, Formal analysis, **CB** : Validation, Supervision, Writing - Review & Editing, **KB** : Conceptualization, Methodology, Validation, Writing - Review & Editing, Visualization, Supervision, Project administration, Funding acquisition.

## References

[1] L. Andriolo, A. Di Martino, S.A. Altamura, A. Boffa, A. Poggi, M. Busacca, S. Zaffagnini, G. Filardo, Matrix-assisted chondrocyte transplantation with bone grafting for knee osteochondritis dissecans: stable results at 12 years, Knee Surg. Sports Traumatol. Arthrosc. 29 (2021) 1830–1840. 10.1007/s00167-020-06230-y.

[2] P.D. Benya, J.D. Shaffer, Dedifferentiated chondrocytes reexpress the differentiated collagen phenotype when cultured in agarose gels, Cell 30 (1982) 215–224. 10.1016/0092-8674(82)90027-7.

[3] L. Duan, B. Ma, Y. Liang, J. Chen, W. Zhu, M. Li, D. Wang, Cytokine networking of chondrocyte dedifferentiation in vitro and its implications for cell-based cartilage therapy, Am J Transl Res 7 (2015) 194–208.

[4] A. Brandl, P. Angele, C. Roll, L. Prantl, R. Kujat, B. Kinner, Influence of the growth factors PDGF-BB, TGF-β1 and bFGF on the replicative aging of human articular chondrocytes during in vitro expansion, Journal Orthopaedic Research 28 (2010) 354–360. 10.1002/jor.21007.

[5] J. Ahn, H. Kumar, B.-H. Cha, S. Park, Y. Arai, I. Han, S.G. Park, S.-H. Lee, AIMP1 downregulation restores chondrogenic characteristics of dedifferentiated/degenerated chondrocytes by enhancing TGF-β signal, Cell Death Dis 7 (2016) e2099–e2099. 10.1038/cddis.2016.17.

[6] S. Speichert, N. Molotkov, K. El Bagdadi, A. Meurer, F. Zaucke, Z. Jenei-Lanzl, Role of Norepinephrine in IL-1β-Induced Chondrocyte Dedifferentiation under Physioxia, IJMS 20 (2019) 1212. 10.3390/ijms20051212.

[7] L. Zhang, A. He, Z. Yin, Z. Yu, X. Luo, W. Liu, W. Zhang, Y. Cao, Y. Liu, G. Zhou, Regeneration of human-ear-shaped cartilage by co-culturing human microtia chondrocytes with BMSCs, Biomaterials 35 (2014) 4878–4887. 10.1016/j.biomaterials.2014.02.043.

[8] X. Huang, Y. Hou, L. Zhong, D. Huang, H. Qian, M. Karperien, W. Chen, Promoted Chondrogenesis of Cocultured Chondrocytes and Mesenchymal Stem Cells under Hypoxia Using In-situ Forming Degradable Hydrogel Scaffolds, Biomacromolecules 19 (2018) 94–102. 10.1021/acs.biomac.7b01271.

[9] Y. Mao, T. Hoffman, A. Wu, J. Kohn, An Innovative Laboratory Procedure to Expand Chondrocytes with Reduced Dedifferentiation, CARTILAGE 9 (2018) 202–211. 10.1177/1947603517746724.

[10] J. Bonaventure, N. Kadhom, L. Cohen-Solal, K.H. Ng, J. Bourguignon, C. Lasselin, P. Freisinger, Reexpression of Cartilage-Specific Genes by Dedifferentiated Human Articular Chondrocytes Cultured in Alginate Beads, Experimental Cell Research 212 (1994) 97–104. 10.1006/excr.1994.1123.

[11] W. Zhang, Y. Zhu, J. Li, Q. Guo, J. Peng, S. Liu, J. Yang, Y. Wang, Cell-Derived Extracellular Matrix: Basic Characteristics and Current Applications in Orthopedic Tissue Engineering, Tissue Engineering Part B: Reviews 22 (2016) 193–207. 10.1089/ten.teb.2015.0290.

[12] A.I. Hoch, V. Mittal, D. Mitra, N. Vollmer, C.A. Zikry, J.K. Leach, Cell-secreted matrices perpetuate the bone-forming phenotype of differentiated mesenchymal stem cells, Biomaterials 74 (2016) 178–187. 10.1016/j.biomaterials.2015.10.003.

[13] J.N. Harvestine, H. Orbay, J.Y. Chen, D.E. Sahar, J.K. Leach, Cell-secreted extracellular matrix, independent of cell source, promotes the osteogenic differentiation of human stromal vascular fraction, J. Mater. Chem. B 6 (2018) 4104–4115. 10.1039/C7TB02787G.

[14] B.S. Kim, J.S. Choi, J.D. Kim, Y.C. Choi, Y.W. Cho, Recellularization of decellularized human adipose-tissue-derived extracellular matrix sheets with other human cell types, Cell Tissue Res 348 (2012) 559–567. 10.1007/s00441-012-1391-y.

[15] J.A. DeQuach, V. Mezzano, A. Miglani, S. Lange, G.M. Keller, F. Sheikh, K.L. Christman, Simple and High Yielding Method for Preparing Tissue Specific Extracellular Matrix Coatings for Cell Culture, PLoS ONE 5 (2010) e13039. 10.1371/journal.pone.0013039.

[16] Y. Mao, T. Block, A. Singh-Varma, A. Sheldrake, R. Leeth, S. Griffey, J. Kohn, Extracellular matrix derived from chondrocytes promotes rapid expansion of human primary chondrocytes in vitro with reduced dedifferentiation, Acta Biomaterialia 85 (2019) 75–83. 10.1016/j.actbio.2018.12.006.

[17] R. Fischer, B.M. Kessler, Gel-aided sample preparation (GASP)—A simplified method for gel-assisted proteomic sample generation from protein extracts and intact cells, Proteomics 15 (2015) 1224–1229. 10.1002/pmic.201400436.

[18] T. Rozario, D.W. DeSimone, The extracellular matrix in development and morphogenesis: A dynamic view, Developmental Biology 341 (2010) 126–140. 10.1016/j.ydbio.2009.10.026.

[19] A.D. Theocharis, S.S. Skandalis, C. Gialeli, N.K. Karamanos, Extracellular matrix structure, Advanced Drug Delivery Reviews 97 (2016) 4–27. 10.1016/j.addr.2015.11.001.

[20] L.E. Niklason, Understanding the Extracellular Matrix to Enhance Stem Cell-Based Tissue Regeneration, Cell Stem Cell 22 (2018) 302–305. 10.1016/j.stem.2018.02.001.

[21] M. Chen, Z. Jiang, X. Zou, X. You, Z. Cai, J. Huang, Advancements in tissue engineering for articular cartilage regeneration, Heliyon 10 (2024) e25400. 10.1016/j.heliyon.2024.e25400.

[22] Y. Wang, M. Du, T. Wu, T. Su, L. Ai, D. Jiang, The application of ECM-derived biomaterials in cartilage tissue engineering, Mechanobiology in Medicine 1 (2023) 100007. 10.1016/j.mbm.2023.100007.

[23] H. Cao, X. Wang, M. Chen, Y. Liu, X. Cui, J. Liang, Q. Wang, Y. Fan, X. Zhang, Childhood Cartilage ECM Enhances the Chondrogenesis of Endogenous Cells and Subchondral Bone Repair of the Unidirectional Collagen–dECM Scaffolds in Combination with Microfracture, ACS Appl. Mater. Interfaces 13 (2021) 57043–57057. 10.1021/acsami.1c19447.

[24] D.S. Harburger, D.A. Calderwood, Integrin signalling at a glance, Journal of Cell Science 122 (2009) 159–163. 10.1242/jcs.018093.

[25] J.D. Humphrey, E.R. Dufresne, M.A. Schwartz, Mechanotransduction and extracellular matrix homeostasis, Nat Rev Mol Cell Biol 15 (2014) 802–812. 10.1038/nrm3896.

[26] E. Charlier, C. Deroyer, F. Ciregia, O. Malaise, S. Neuville, Z. Plener, M. Malaise, D. De Seny, Chondrocyte dedifferentiation and osteoarthritis (OA), Biochemical Pharmacology 165 (2019) 49–65. 10.1016/j.bcp.2019.02.036.

[27] Y. Chen, Y. Yu, Y. Wen, J. Chen, J. Lin, Z. Sheng, W. Zhou, H. Sun, C. An, J. Chen, W. Wu, C. Teng, W. Wei, H. Ouyang, A high-resolution route map reveals distinct stages of chondrocyte dedifferentiation for cartilage regeneration, Bone Res 10 (2022) 38. 10.1038/s41413-022-00209-w.

[28] W. Su, Y. Nie, S. Zheng, Y. Yao, Recent Research on Chondrocyte Dedifferentiation and Insights for Regenerative Medicine, Biotech & Bioengineering 122 (2025) 749–760. 10.1002/bit.28915.

[29] E.S. Place, N.D. Evans, M.M. Stevens, Complexity in biomaterials for tissue engineering, Nature Mater 8 (2009) 457–470. 10.1038/nmat2441.

[30] W.P. Daley, K.M. Yamada, ECM-modulated cellular dynamics as a driving force for tissue morphogenesis, Current Opinion in Genetics & Development 23 (2013) 408–414. 10.1016/j.gde.2013.05.005.

[31] C. Liu, M. Pei, Q. Li, Y. Zhang, Decellularized extracellular matrix mediates tissue construction and regeneration, Front Med 16 (2022) 56–82. 10.1007/s11684-021-0900-3.

[32] S. Aghlara-Fotovat, A. Nash, B. Kim, R. Krencik, O. Veiseh, Targeting the extracellular matrix for immunomodulation: applications in drug delivery and cell therapies, Drug Deliv. and Transl. Res. 11 (2021) 2394–2413. 10.1007/s13346-021-01018-0.

[33] J.K. Kular, S. Basu, R.I. Sharma, The extracellular matrix: Structure, composition, age-related differences, tools for analysis and applications for tissue engineering, J Tissue Eng 5 (2014) 2041731414557112. 10.1177/2041731414557112.

[34] J.L. Dziki, D.S. Wang, C. Pineda, B.M. Sicari, T. Rausch, S.F. Badylak, Solubilized extracellular matrix bioscaffolds derived from diverse source tissues differentially influence macrophage phenotype, J Biomedical Materials Res 105 (2017) 138–147. 10.1002/jbm.a.35894.

[35] I.T. Swinehart, S.F. Badylak, Extracellular matrix bioscaffolds in tissue remodeling and morphogenesis, Developmental Dynamics 245 (2016) 351–360. 10.1002/dvdy.24379.

[36] M. Laude, V. Kolliopoulos, A.G. Mikos, L.J. White, E. Cosgriff-Hernandez, Extracellular-Matrix-Based Materials from Decellularized Tissue: Opportunities, Challenges, and Future Directions in Regenerative Medicine, Adv Healthcare Materials 15 (2026) e02107. 10.1002/adhm.202502107.

[37] Y.A. Kadry, D.A. Calderwood, Chapter 22: Structural and signaling functions of integrins, Biochimica et Biophysica Acta (BBA) - Biomembranes 1862 (2020) 183206. 10.1016/j.bbamem.2020.183206.

[38] S. Dupont, L. Morsut, M. Aragona, E. Enzo, S. Giulitti, M. Cordenonsi, F. Zanconato, J. Le Digabel, M. Forcato, S. Bicciato, N. Elvassore, S. Piccolo, Role of YAP/TAZ in mechanotransduction, Nature 474 (2011) 179–183. 10.1038/nature10137.

[39] Z. Xie, M. Khair, I. Shaukat, P. Netter, D. Mainard, L. Barré, M. Ouzzine, Non-canonical Wnt induces chondrocyte de-differentiation through Frizzled 6 and DVL-2/B-raf/CaMKIIα/syndecan 4 axis, Cell Death Differ 25 (2018) 1442–1456. 10.1038/s41418-017-0050-y.

[40] K. Katoh, Integrin and Its Associated Proteins as a Mediator for Mechano-Signal Transduction, Biomolecules 15 (2025) 166. 10.3390/biom15020166.

[41] P. Kanchanawong, D.A. Calderwood, Organization, dynamics and mechanoregulation of integrin-mediated cell–ECM adhesions, Nat Rev Mol Cell Biol 24 (2023) 142–161. 10.1038/s41580-022-00531-5.

[42] E. Silk, H. Zhao, H. Weng, D. Ma, The role of extracellular histone in organ injury, Cell Death Dis 8 (2017) e2812–e2812. 10.1038/cddis.2017.52.

[43] B. Singh, M. Fredriksson Sundbom, U. Muthukrishnan, B. Natarajan, S. Stransky, A. Görgens, J.Z. Nordin, O.P.B. Wiklander, L. Sandblad, S. Sidoli, S. El Andaloussi, M. Haney, J.D. Gilthorpe, Extracellular Histones as Exosome Membrane Proteins Regulated by Cell Stress, J of Extracellular Vesicle 14 (2025) e70042. 10.1002/jev2.70042.

